# The trajectory of discrete gating charges in a voltage-gated potassium channel

**DOI:** 10.1101/2020.04.23.058818

**Authors:** Michael F. Priest, Elizabeth E.L. Lee, Francisco Bezanilla

## Abstract

Positively-charged amino acids respond to membrane potential changes to drive voltage sensor movement in voltage-gated ion channels, but determining the trajectory of voltage sensor gating charges has proven difficult. We optically tracked the movement of the two most extracellular charged residues (R1, R2) in the Shaker potassium channel voltage sensor using a fluorescent positively-charged bimane derivative (qBBr) that is strongly quenched by tryptophan. By individually mutating residues to tryptophan within the putative trajectory of gating charges, we observed that the charge pathway during activation is a rotation and a tilted translation that differs between R1 and R2 and is distinct from their deactivation pathway. Tryptophan-induced quenching of qBBr also indicates that a crucial residue of the hydrophobic plug is linked to the Cole-Moore shift through its interaction with R1. Finally, we show that this approach extends to additional voltage-sensing membrane proteins using the *Ciona intestinalis* voltage sensitive phosphatase (CiVSP).

## Introduction

The nature of the motion of the voltage sensor in voltage-gated ion channels has been a subject of intensive research. This motion is driven by voltage changes sensed by positively-charged amino acids (typically arginines) found on the fourth transmembrane segment (S4) of each monomer of the tetrameric channel. The number of charged amino acids driving this motion varies from channel to channel, but has been shown in the canonical voltage-gated Shaker potassium channel (Kv) to consist of the four most extracellular arginines (Aggarwal & MacKinnon, 1996; Seoh et al., 1996). However, research into the motion of these individual gating charges has suffered from experimental limitations. Replacement of arginine by histidine together with pH titration allows the study of gating charge end positions during activation, but does not show the charge’s actual trajectory (Starace et al., 1997; Starace & Bezanilla, 2001). Cross-linking studies replace residues with cysteines to see which residues interact with each other in different conformational states, necessarily abrogating the positive charge of any gating charge if they are examined directly (Broomand et al., 2003; Henrion et al., 2012; Lainé et al., 2003), and accessibility studies have the same limitation (Yang & Horn, 1995). Similarly, site directed fluorimetric approaches typically replace a residue with a cysteine and then attach a fluorescent dye, providing the additional advantage of being able to monitor conformational changes in real-time, but do not directly follow the movement of the gating charges (Cha & Bezanilla, 1997; Mannuzzu et al., 1996; Priest & Bezanilla, 2015). Positively charged adducts such as methanethiosulfonate-ethyltrimethylammonium (MTSET) linked to cysteines placed at gating charges can be used to replace the positive charge and characterize discrete gating charges (Ahern & Horn, 2004, 2005; Baker et al., 1998; Larsson et al., 1996). However, these replacement charges cannot be rapidly monitored, limiting their use for observing conformational changes.

Ideally, one would like to follow the movement of individual gating charges in real time as they respond to changes in the electric field. This requires a fluorophore that is comparable to the gating arginines. We used monobromo(trimethylammonio)bimane (qBBr), a small molecule fluorescent dye with a permanent positive charge (Figure 1A). While bulkier than an arginine (MW≈295 versus 101) (Figure 1B), modeling the conjugation of qBBr to a cysteine substituted into the two most extracellular gating charges in a Shaker-type voltage-gated potassium channel (R362 or R1 and R365 or R2) (Chen et al., 2010; Pettersen et al., 2004) provided a distance between the carbon backbone and the positive charge of qBBr of ∼5.5 Å, very close to the analogous distance of ∼5.3 to 6.5 Å for arginine. Therefore, qBBr attached to a cysteine at the endogenous position of the gating charge may mimic the positive gating charge. In addition to being positively charged, qBBr has useful fluorescence quenching properties. Tryptophan has been shown to strongly quench qBBr fluorescence, with weak quenching by tyrosine, and no quenching from various other amino acids including histidine, phenylalanine, methionine, aspartate, or arginine (Mansoor et al., 2010). This phenomenon of ‘tryptophan-induced quenching’ in bimane dyes generally (Mansoor et al., 2002), together with their remarkable environmental insensitivity (Mansoor et al., 1999), has been taken advantage of to measure conformational rearrangements of various membrane proteins, including the β2-adrenergic G protein-coupled receptor (Yao et al., 2006), a cyclic nucleotide-gated ion channel (Islas & Zagotta, 2006), the BK channel (Semenova et al., 2009), a proton-gated ion channel (Menny et al., 2017), and a lactose permease (Smirnova et al., 2014). In general, these studies use the fluorescent quenching produced by an interaction between bimane and a particular tryptophan, whether native or engineered into the protein, to provide insight into how the protein moves during conformational transitions.

**Figure 1.**
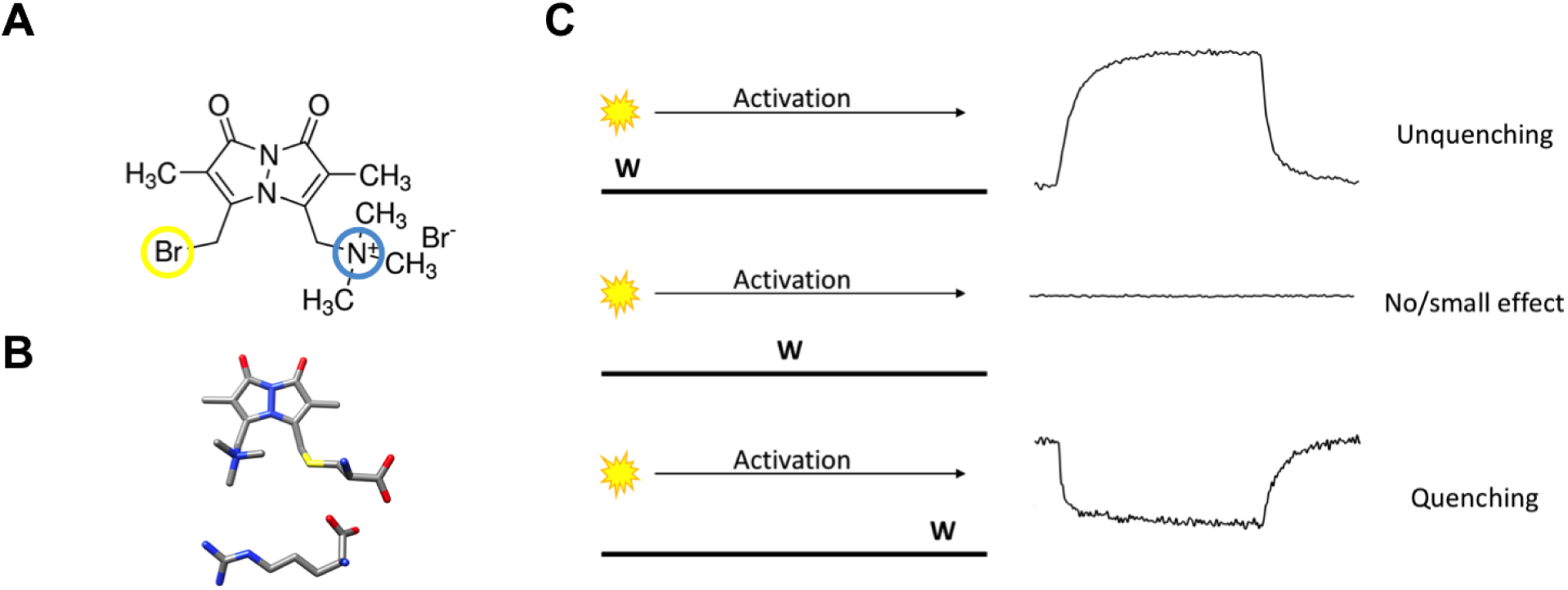
The basic principles of qBBr gating charge tracking in time. (A) Chemical structure of monobromo(trimethylammonio)bimane, a small positively charged (blue) fluorophore with the ability to conjugate to cysteine (yellow). (B) Size comparison of qBBr-Cys (top) to an arginine (bottom). (Blue, nitrogen; red, oxygen; sulfur, yellow; grey, carbon). (C) Cartoon schematic of qBBr-tryptophan distance-based quenching with representative fluorescence data for unquenching (top), no effect (center), and quenching (bottom).

Here, we perform site-directed voltage-clamp fluorimetry on qBBr-bound Shaker Kv channels expressed in *Xenopus laevis* oocytes. By substituting a cysteine for a native gating charge and then covalently attaching qBBr to this site, we produce a fluorescent mimic of a discrete charge in the voltage sensor of the channel. Individual native or engineered tryptophans in the channel are then used to quench the fluorescence of the charged qBBr as it moves in response to changes in membrane potential. We used this system to investigate the pathway taken by R1 and R2 during activation and deactivation, as well as to examine the effect of the relaxed state (Lacroix et al., 2011; Villalba-Galea et al., 2008) on these individual gating charges. In general, the movement of R1C- and R2C-qBBr follow a tilted translation across the membrane and rotation, giving new information on the trajectories that have been inferred from recent consensus models on the motion of the voltage sensor (Li et al., 2014; Vargas et al., 2011, 2012). Of particular interest, during normal channel activation R1C-qBBr does not appear to interact with F290W, the most intracellular residue of the hydrophobic plug (Chen et al., 2010; Lacroix & Bezanilla, 2011), also called the gating charge transfer center (Tao et al., 2010). Instead, R1 only interacts with F290W at extremely negative potentials, suggesting that R1 does not normally move past the hydrophobic plug and provides the basis of the Cole-Moore shift. This technique should also be transferrable to other voltage-sensing membrane proteins; as a proof of principle, we demonstrate its use in the voltage-sensitive phosphatase CiVSP.

## Results

### The basic principles of qBBr gating charge tracking in time

The idea of the present approach is to study the translocation of the gating charge (now a fluorophore, qBBr) as the membrane potential is changed using the specific quenching of qBBr by a tryptophan (W) that is positioned nearby or in the path of qBBr.

If we were tracking fluorescence at the single molecule level, the fluorescence signal we would observe would depend on the position of the W with respect to the moving qBBr. Let us assume that we are applying a positive voltage to activate the voltage sensor that moves between two discrete positions. We can distinguish three extreme cases schematically (Figure 1C, left panel). If the W is near the resting position of the qBBr, we would see a sudden increase in fluorescence when the qBBr moves away from it and the time lag before that increase corresponds to the waiting time of the sensor before it jumps across the energy barrier (Figure 1C, left, top trace). On the other hand, if the W is in the path of qBBr (Figure 1C, left, middle), we would see an extremely brief decrease of fluorescence as the qBBr passes by the quenching group. Finally, if the W is near the final position of the qBBr (Figure 1C, left, bottom) we would see a sudden decrease in fluorescence that would be maintained when the qBBr reaches that point. The duration of the high fluorescence period is the waiting time before the sensor crosses the energy barrier.

However, since we are looking at a large ensemble of molecules the macroscopic fluorescence signals we see are the result of an ensemble of voltage-sensing domains (VSDs) moving, each one at a different time according to its first latency duration, generating a continuous change in fluorescence. The fast event for the case of a W in the middle of the qBBr path would not be visible in the ensemble because the crossing of the VSDs is unsynchronized and if it is exactly in the middle no signal will be seen (Figure 1C, right panel, middle trace). The other two extreme cases will generate a continuous quenching or unquenching depending on the placement of the tryptophan (Figure 1C, right, top and bottom).

A simple model of quenching of qBBr by W is

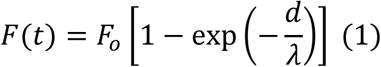

where *F(t)* is qBBr fluorescence, *F*_*0*_ is the maximum qBBr fluorescence (no quenching), *d* is the distance from qBBr to W and ⍰ is the characteristic distance for quenching, Now, we assume that qBBr moves in a trajectory represented by *x* between the initial position, x_0_, and the final position, x_f_, and W is located somewhere near the x trajectory, x_W_. Let us consider first that qBBr moves between two states and define *P*_*0*_*(t)* and *P*_*f*_*(t)* as the probabilities that qBBr is in *x*_*0*_ and *x*_*f*_ respectively. Then, *P*_*0*_*(t) + P*_*f*_*(t)=*1 and we can write the time course of the fluorescence as

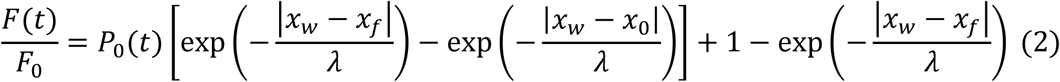

This means that the time course of fluorescence will have the time course of *P*_*0*_*(t)* and will not depend on the position of W in the trajectory.

If we now assume that the number of states is more than 2, say i=1, 2, 3, …, n, we can write a general solution for the fluorescence

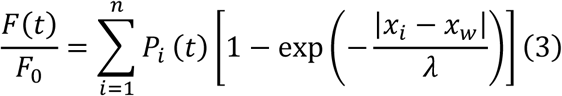

*P*_*i*_*(t)* will have *n-1* eigenvalues, so the time course of fluorescence will now depend on the position of the W in the trajectory, because *P*_*i*_*(t)* will be weighted by the exponential term that includes the distance to each position. Simulations with 3 states show fluorescence time courses with more than one exponential component and, depending on the position and dwell time in the intermediate state, the response may be biphasic (i.e. fluorescence increases and decreases) during the depolarization. This is expected because as it traverses its trajectory the qBBr will go through periods when it will be closer to the W and others when it will be further away. This is a prediction that will be important in interpreting the experimental results.

### qBBr mimics a native gating charge

While qBBr crucially contains a permanent charge (Figure 2A) like the arginines that constitute the gating charges in the Shaker Kv channel, this is not a guarantee that the voltage sensor will continue to behave normally with a fluorescent dye substituted for a gating charge. To test whether this was the case, we first separately mutated R1 and R2 in a non-conducting (W434F), non-fast inactivating (Δ6-46) Shaker channel to cysteines. Following expression of these constructs in *Xenopus laevis* oocytes, qBBr was conjugated to either of these charges by incubation of the oocytes in a depolarizing solution containing qBBr. Oocytes containing qBBr conjugated to R1C or R2C were then simultaneously recorded optically and electrophysiologically using the cut-open oocyte voltage clamp technique. In both cases, the observed gating currents are likely a combination of voltage sensors that have been labeled with qBBr and unlabeled voltage sensors that retain a bare cysteine. For R1C-qBBr, gating currents appeared to be faster than those of the wild type channel but still comparable in voltage sensitivity (Figures 2A and 2B). With R2C-qBBr, gating currents were also more rapid, with voltage sensor movement beginning at more hyperpolarized potentials than in the wild-type channel and producing a shallow Q-V curve (Figures 2A and 2B). These findings suggest that the additional bulk of qBBr may destabilize the resting state of the voltage sensor when it is conjugated to the more intracellular R2C, but not when it is conjugated to the more extracellular R1C. This is in good agreement with numerous computational models of the resting state of the Kv channel that suggest that R1 is less sterically inhibited in the resting state than R2 (Vargas et al., 2012). They also suggest that qBBr would act as a faithful mimic of the movement of the gating charge of R1, and should also mimic the movement of R2, albeit with energetic differences.

**Figure 2.**
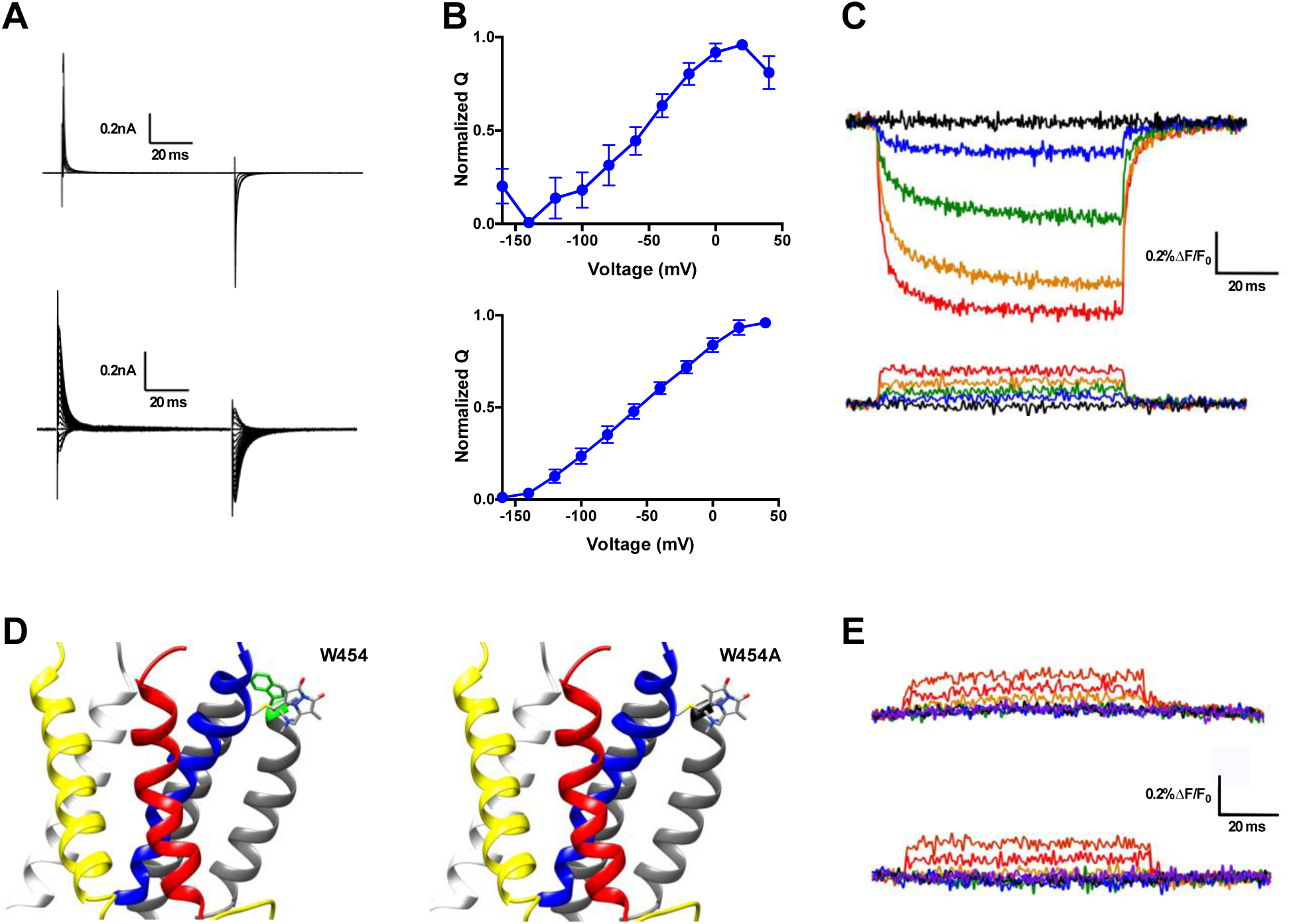
qBBr mimics a native gating charge. (A) Representative gating current traces of Shaker R1C-qBBr (top) and R2C-qBBr (bottom). (B) Normalized charge (Q) versus voltage (V) QV curves of R1C-qBBr (top, n=4) and R2C-qBBr (bottom, n=3). Data are shown as mean ± SEM. (C) Representative fluorescence traces of qBBr labeled R1C-qBBr (top) and R2C-qBBr (bottom). In all figures, membrane potentials during the pulse are: brown, +80 mV; red, +40 mV; orange, 0 mV; green, −40 mV; blue, −80 mV; black, −120 mV; purple, −160 mV. (D) Structures showing the position of the in-silico mutation 454 from a tryptophan (left) to an alanine (right) (PDB: 3LUT). In all Shaker structures, transmembrane domains (S1-S6) are colored: S1, white; S2, yellow; S3, red; S4, blue; S5 and S6, grey. (E) Representative fluorescence traces for R1C-qBBr:W454A (top) and R1C-qBBr:W454F (bottom).

Upon depolarization, the fluorescence signal of R1C-qBBr decreased dramatically (Figure 2C, top), while the corresponding fluorescence signal of R2C-qBBr did not (Figure 2C, bottom). A reduction in qBBr fluorescence is likely due to the presence of a tryptophan, or possibly a tyrosine. Since the fluorescence signal is lower when the R1C-qBBr is in the active state than when it is in the resting state, the quenching residue should be near the active state of R1. Based on a homology model of a Kv channel crystal structure in the relaxed state (Chen et al., 2010), R1 should reside near an endogenous tryptophan (W454) at the extracellular side of S6 (Figure 2D). Additionally, it has been reported that R1 comes into close proximity to this region of the channel (Lainé et al., 2003). Mutation of this tryptophan to a non-quenching alanine abolished the fluorescence reduction that we observed during activation (Figure 2E, top). The mutation of this tryptophan to a phenylalanine also abolished the endogenous voltage-dependent fluorescence (Figure 2E, bottom). However, as this mutation reduced expression, we proceeded with a W454A background. Therefore, our optical data supports the idea that upon activation, R1C-qBBr comes near the pore domain where its fluorescence is quenched by W454, while R2C-qBBr does not. These results are in agreement with crystallographic and functional data and suggest that R2C-qBBr and R1C-qBBr may mimic the motion of the native gating charges.

### An exogenously substituted tryptophan (E247W) produces voltage-dependent fluorescence changes in R1C-qBBr

To map the trajectory of R1 and R2 using qBBr, we created a series of constructs based on the R1C:W454A and the R2C backgrounds. We used these two constructs as backgrounds because they only showed a very small depolarization-induced increase in fluorescence that had a fast time course with no voltage dependent kinetics (Figure 2E) which we were unable to link to any endogenous tyrosine or tryptophan (Figure 2–figure supplement 1). As this residual fluorescence change was unobservable in the presence of tryptophan-induced quenching in the R1C-qBBr:W454W construct, we hypothesized that substituting a tryptophan elsewhere into the channel would similarly alter the qBBr fluorescence signal in a way that would dominate over the residual fluorescence signal.

As an example, we substituted a tryptophan at position 247 of the R1C-W454A construct, generating R1C-qBBr:W454A;E247W (Fig. 3A). In response to changes in membrane potential, the exogenous, substituted tryptophan produced voltage-induced fluorescence changes (Figure 3B) that were markedly different from the background fluorescence changes of R1C-qBBr:W454A (Figure 2E, top). If R1C-qBBr is closer to the tryptophan at 247 in the resting state than in the active state, we should observe an increase in fluorescence upon activation (Figure 1C, top).The experiment shows that during activation, E247W produces a clear unquenching, or increase in fluorescence, of R1C-qBBr (Figure 3B). Therefore, R1C-qBBr moves away from position 247 upon activation.

**Figure 3.**
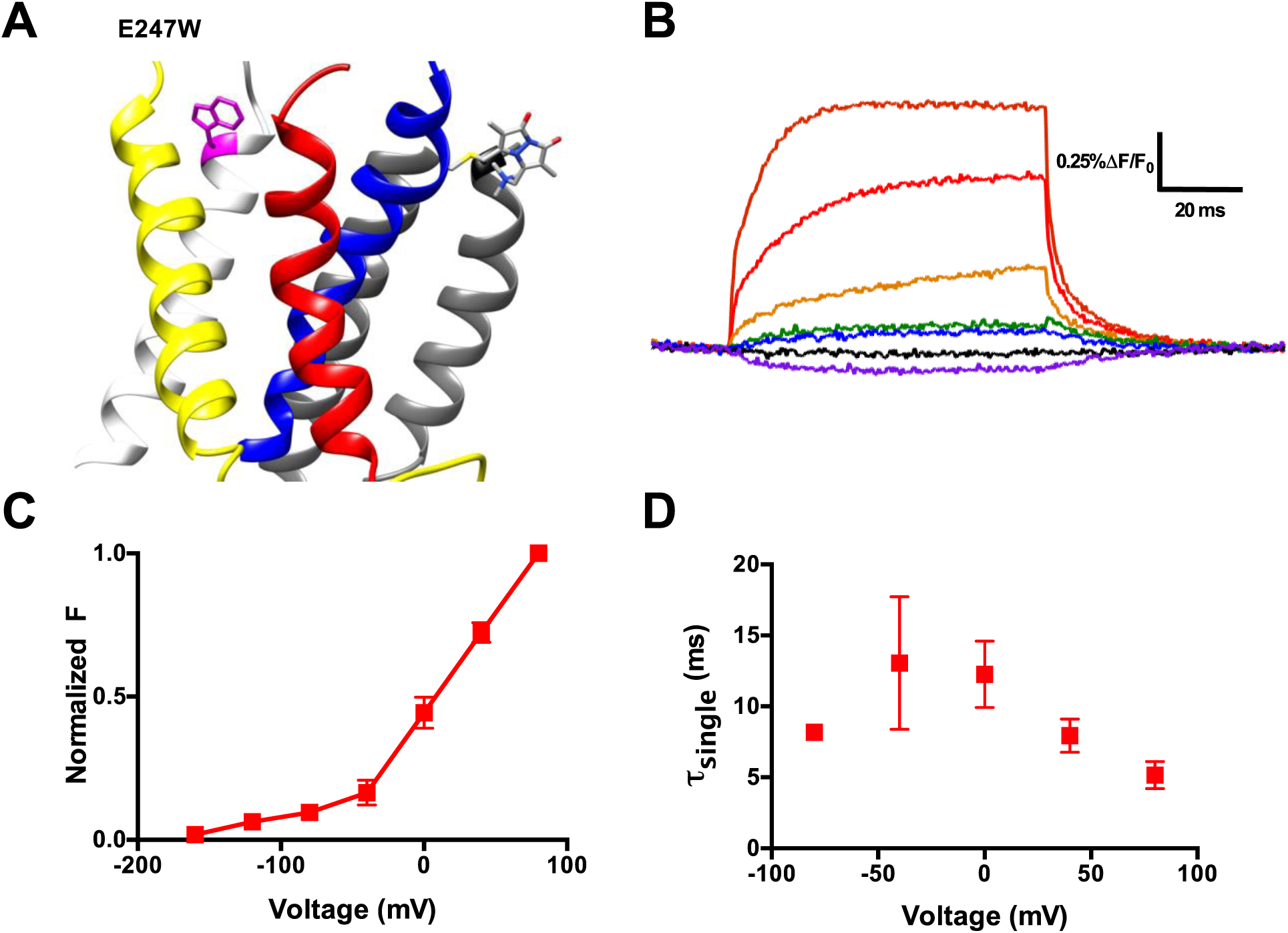
An exogenously substituted tryptophan (E247W) produces voltage-dependent fluorescence changes in R1C-qBBr. (A) Homology structure of Shaker construct, R1C-qBBr:W454A;E247W demonstrating the tryptophan placement within the protein. (B) A representative activation family of fluorescence traces. (C) A normalized fluorescence vs voltage curve. (D) A single exponential **τ** of fluorescence upon activation (n=3). Data are shown as mean ± SEM.

Examining the fluorescence signal more closely, we see that it has many properties reflecting movement of the voltage sensor. Just as the gating currents produced by the voltage sensor are voltage-sensitive, the fluorescence signal of R1C-qBBr:W454A;E247W has a voltage-sensitive response. From our family of fluorescence traces (Figure 3B) we can generate a change in fluorescence versus voltage (FV) curve and observe the amount of fluorescence quenching or unquenching due to changes in voltage (Figure 3C). Moreover, the voltage-sensitivity of the fluorescence is also seen in the kinetics of the fluorescence traces (Fig. 3D).

### Activation pathways mapped by qBBr and substituted tryptophan-induced fluorescence quenching

We generated numerous constructs with an exogenous, substituted tryptophan that produced voltage-induced fluorescence changes in R1C-qBBr:W454A or R2C-qBBr that were distinct from the background fluorescence changes and from each other (Figures 3B, 3C, 4A and 4B). A decrease in fluorescence upon depolarization suggests that the R1C-or R2C-qBBr comes closer to the inserted tryptophan residue during the transition from the resting to the active state, while an increase in fluorescence upon depolarization suggests that the gating charge moves away from that residue during activation (Figure 1C). Therefore, by pooling the data from numerous constructs, each with a single tryptophan, we can map the activation pathways of R1 and R2 (Figures 4A and 5A).

**Figure 4.**
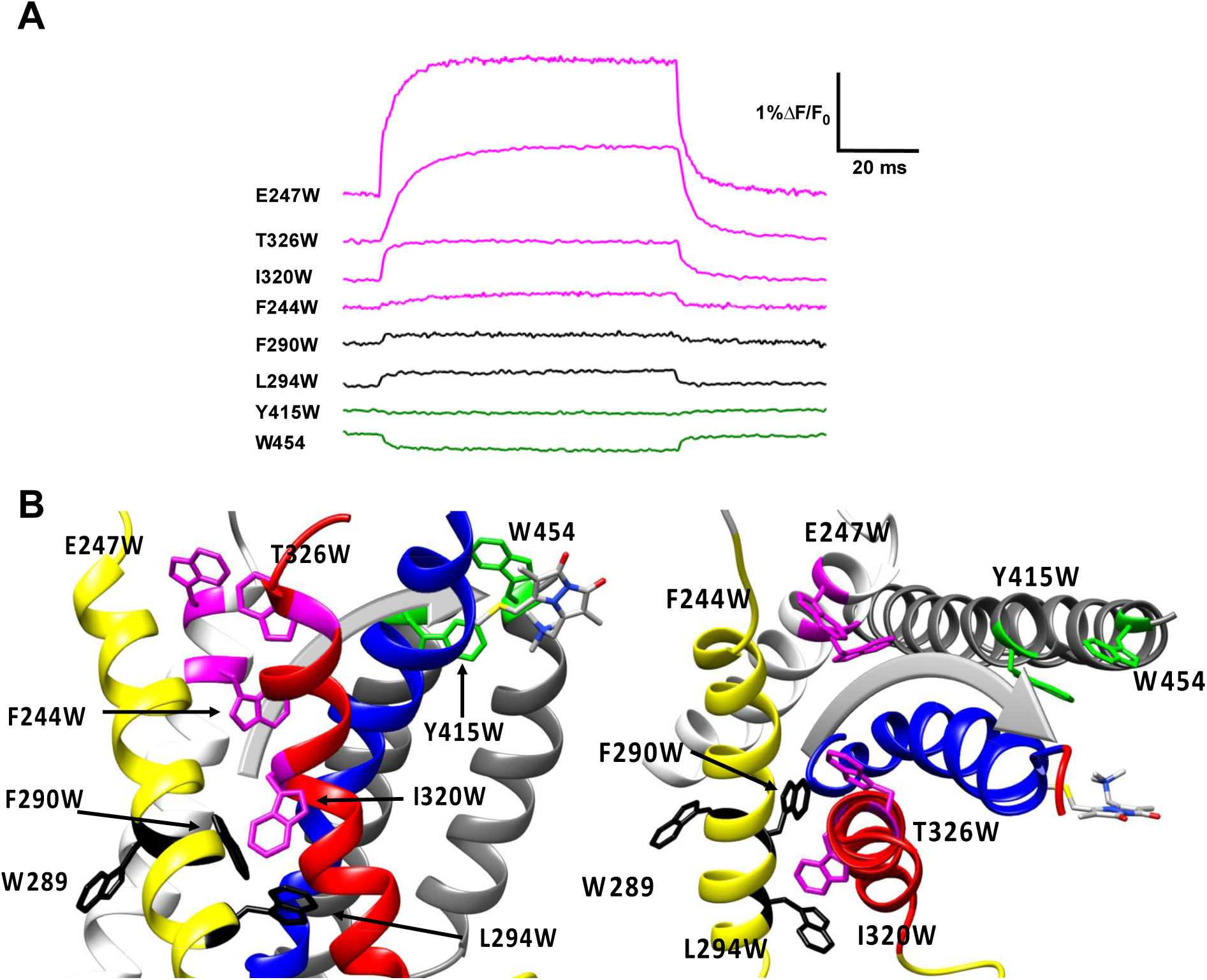
Activation pathway of R1C-qBBr:W454A mapped by several individually substituted tryptophans. (A) Representative qBBr traces at +80mV. As qBBr moves closer to a W, the W quenches (green) the qBBr fluorescence (Y415W, W454). When it moves further away from a W, the qBBr fluorescence is unquenched (pink; F244W, E247W, I320W, T326W). Some W mutations have no effect (black; 289W, L294W, F290W). (B) A summary structure of activation of R1C-qBBr, with side (left) and extracellular (right) views of the VSD. The activation pathway based on the W quenching/unquenching for R1C-qBBr is marked by a grey arrow.

In response to a depolarizing pulse, R1C-qBBr fluorescence was quenched by the endogenous tryptophan at residue 454, as well as by a tryptophan substituted for the phenylalanine at the extracellular side of the S5 at position 416. R1C-qBBr moved away from tryptophans inserted at F244W, E247W, 320W, and T326W (Figures 4A and 4B). Interestingly, we found no appreciable changes in the fluorescence signal of F290W or L294W in response to depolarizing pulses (Figures 4A and 4B). The small fast fluorescence signal seen for these two constructs is reminiscent of that of R1C-qBBr:W454A, (Figure 2E and Figure 2—figure supplement 1), i.e., the residual fluorescence signal that is small and has fast voltage-independent kinetics. Together these findings map out a pathway for R1 that consists of both an intracellular to extracellular translation, tilted by about 30° with respect to the normal of the membrane plane, and a rotation (Fig. 4B). Specifically, R1C-qBBr seems to reside near the pore domain in the active state, as observed in the crystal structure (Chen et al., 2010). It reaches this position from a resting position that seems to reside approximately one helix turn extracellular to F290, as it shows movement away from I320W, but not from W289W, F290W, or L294W.

Substitution of W residues along the pathway of R2 reveals that upon activation R2C-qBBr moves closer to Y415W, F416W, and T326W, while moving away from I241W, F244W, I287W, and L294W (Figures 5A and B). Therefore, in the active state, R2C-qBBr resides between the extracellular side of the S3 and S5 segments, as quenching is observed from T326W, Y415W, and F416W. As opposed to R1C-qBBr, R2C-qBBr does not encounter the S6 segment upon activation. In the resting state, the fluorescence results suggest that R2C-qBBr is stable near F290, unquenching from a cluster of tryptophans inserted at L294, I287, and I241. As seen with R1, there is a rotation in the movement of R2; however, the rotation of R2 is less evident.

**Figure 5.**
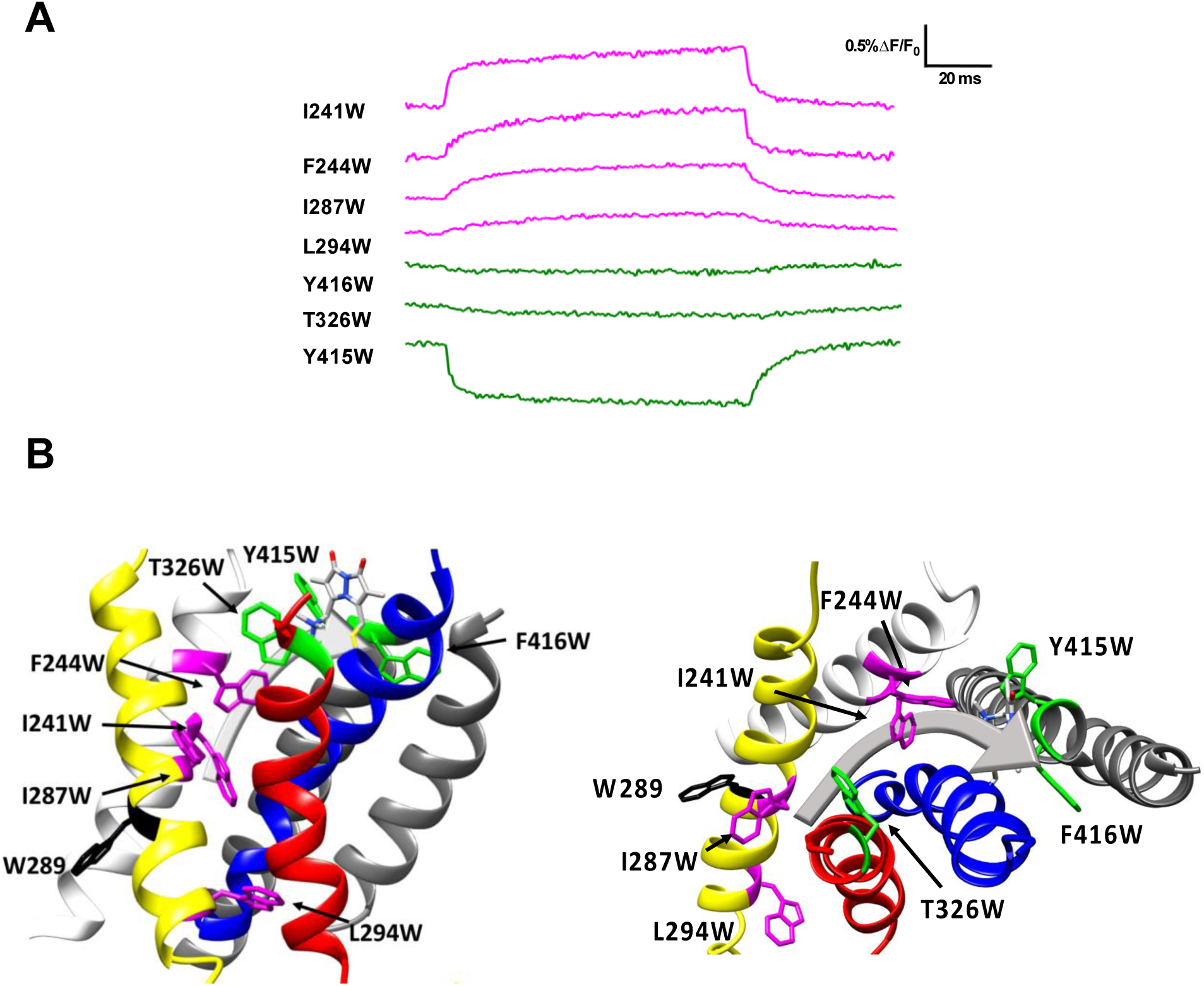
Activation pathway of R2C-qBBr mapped by several individually substituted tryptophans. (A) Representative qBBr traces at +80 mV. As qBBr moves closer to a W, the W quenches (green) the qBBr fluorescence (Y415W, T326W, Y416W). When it moves further away from a W, the qBBr fluorescence is unquenched (pink; I241W, F244W, I287W, L294W). Residue W289 (black) does not affect the fluorescence. (B) A summary structure of activation of R2C-qBBr with side (left) and extracellular (right) views of the VSD. The activation pathway based on the W quenching/unquenching for R2C-qBBr is marked by a grey arrow.

Interestingly, the fluorescence responses differ qualitatively between the two gating charges for tryptophans substituted at L294, T326, and W454. Thus, qBBr mapping reveals that R1 and R2 travel distinct paths in relationship to the voltage-sensing domain during activation. Notable exceptions to this include Y415, W289, and F244, which produce similar responses from both R1C and R2C-qBBr. As it faces the lipid membrane, it is unsurprising that W289 does not quench either R1C or R2C-qBBr; however, this suggests that the S2 in which it resides is unlikely to undergo any large rotations that would expose this residue to the gating charges. Y415W quenches both R1 and R2, underscoring the activated position of both these residues near the extracellular surface of the membrane. The neighboring residue F416W quenched R2C-qBBr and did not express with R1C-qBBr. F416 has previously been shown to bridge to R1 (Conti et al., 2016; Lainé et al., 2003; Phillips & Swartz, 2010) and R2 (Conti et al., 2016). Finally, F244 has been shown to interact with both R1 and R2 (Lacroix et al., 2012), and has been proposed to act as a critical stabilizer of the active state of the voltage sensor (Lacroix et al., 2014). Together, our fluorescence data show that upon activation both R1 and R2 undergo a tilted translation from the intracellular to the extracellular side of the membrane, together with a rotation (Figures 4B and 5B). The discrete and unique pathways mapped out by these charges provide new levels of clarity into how these charges move distinctly from each other.

### The VSD deactivation path differs from that of activation

Previous studies have discussed the asymmetry in activation and deactivation currents, with deactivation currents being slower than those of activation (Labro et al., 2012; Lacroix et al., 2011; Lacroix & Bezanilla, 2012). Specifically, deactivation currents are slowed down in a voltage-dependent manner and correspond with pore opening (McCormack et al., 1994; Perozo et al., 1993). If the path of deactivation was the same as that of activation, we would expect qBBr fluorescence to display similar voltage dependence and kinetics during both activation (Figure 6A, left) and deactivation (Figure 6A, right). The comparison of the activation charge versus voltage curves (QVs) and FVs to the deactivation QVs and FVs shows that during deactivation there are leftward shifts in several R1C-qBBr:W454A constructs: E247W, I320W, and T326W (Figures 6B and C). The shifts in QV and FV curves indicate that the VSD has a different energetic path from the active to the resting state than from the resting state to the active state. This is also reflected in the speed of the fluorescence traces, where we do not see exact overlap in the kinetics of activation and deactivation fluorescence (Figure 6D). In two of our constructs, R1C-qBBr:W454A;E247W and R1C-qBBr:W454A;I320W, the kinetics of the deactivation fluorescence are slower than that of activation. However, we detect the opposite for the fluorescence kinetics of R1C-qBBr:W454A;T326W (Figure 6D, right) where activation fluorescence kinetics are slower than those of deactivation. Others have observed that with the T326W mutation, both activation and deactivation are slowed, but activation much more strongly (Hong & Miller, 2000). Thus, we have strong evidence through both the differing kinetics and the leftward shifts in the FVs of R1C-qBBr fluorescence that a discrete gating charge takes a different path in activation than during deactivation; this is also seen with R2C-qBBr (Figure 6—figure supplement 1).

**Figure 6.**
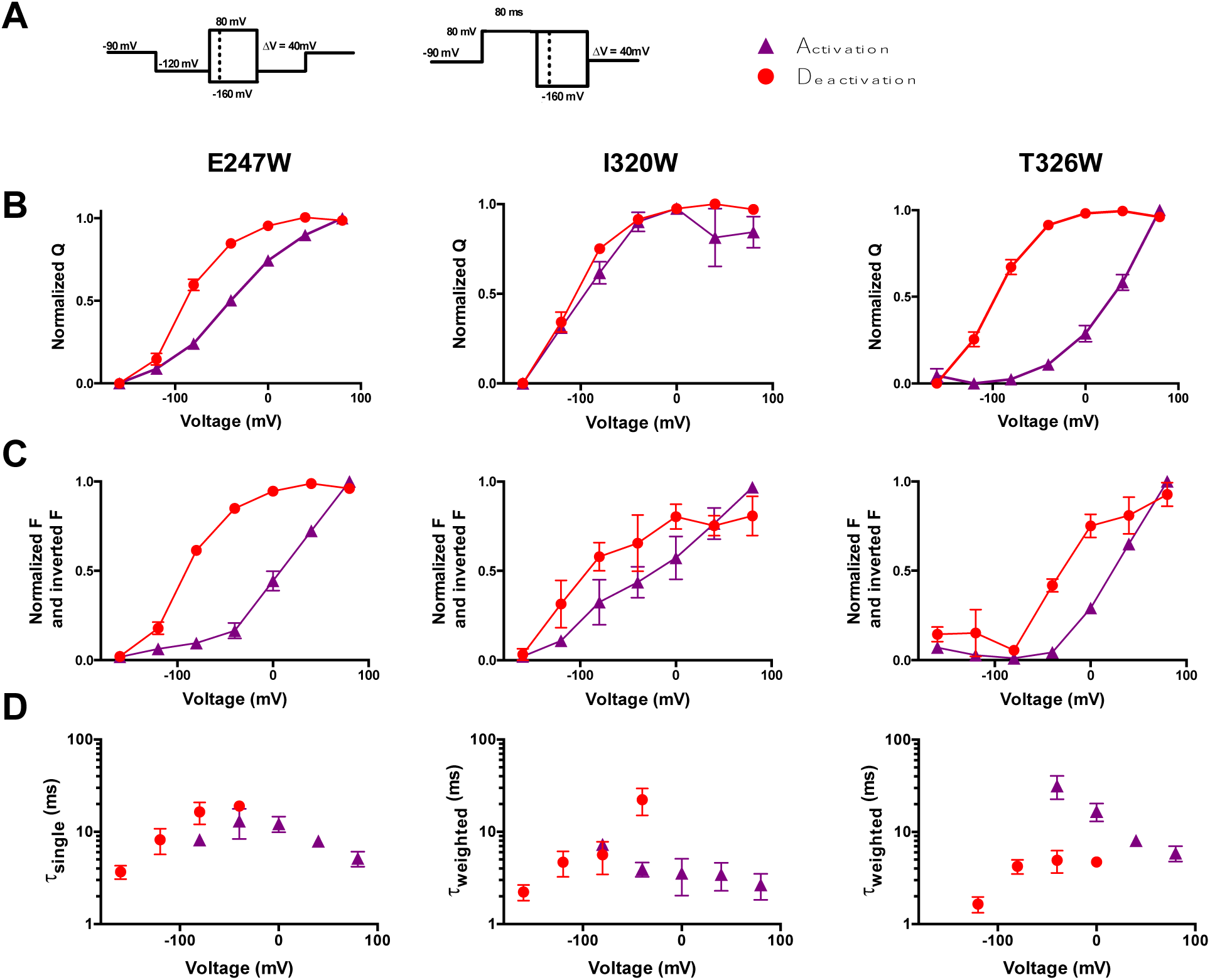
The VSD deactivation path differs from that of activation. (A) Activation (purple) and deactivation (red) protocols. Shading indicates where gating and fluorescence were measured for (B-D). (B) Comparison of activation (purple triangles) and deactivation (red circles) QVs for R1C-qBBr:W454A;E247W (left, n=8), R1C-qBBr:W454A;I320W (center, n=5), and R1C-qBBr:W454A;T326W (right, n=7). (C) As in (B), but a comparison of activation and deactivation FVs, rather than QVs. (D) Comparison of single exponential τ_act_ (red circles, n=3) to τ_dea_ (purple triangles, n=4) of R1C-qBBr:W454A;E247W (left). Weighted τ_act_ (red circles, n=3) to τ_dea_ (purple triangles, n=5) of R1C-qBBr:W454A;I320W (center). Weighted τ_act_ (red circles, n=3) to τ_dea_ (purple triangles, n=3) of R1C-qBBr:W454A;T326W (right). Data are shown as mean ± SEM.

In addition to a resting state and an active state, the voltage sensor has a third state, called the relaxed state, which it enters after prolonged depolarization (F. Bezanilla et al., 1982; Lacroix et al., 2011; Villalba-Galea et al., 2008). This state has been observed in most S4-based voltage sensors, including voltage-gated sodium channels (Nav), Kv, hyperpolarization-activated cyclic nucleotide-gated channels (HCN), and CiVSP (F. Bezanilla et al., 1982; Bruening-Wright & Larsson, 2007; Villalba-Galea et al., 2008). Although there is no charge moved during the transition from the active state to the relaxed state, the relaxed state can be detected through a slowing of the gating kinetics as well as a left-shifted QV curve in R1C-qBBr:W454A;E247W and R1C-qBBr:W454A;T326W (Figure 6—figure supplement 2). As with deactivation, qBBr fluorescence can be used to visualize the relaxed state.

### R1C-qBBr interaction with F290 provides the basis of the Cole-Moore shift

In addition to uncovering information about transitions to and from the resting, active, and relaxed states, qBBr tracking provides novel insight into the Cole-Moore shift. The Cole-Moore shift describes a phenomenon where following a prolonged hyperpolarizing prepulse, a depolarizing pulse elicits a conductance that has a longer time lag, or delay before beginning, than when there is no prepulse; this lag prolongs as the prepulse is made longer and more negative (Cole & Moore, 1960; Hoshi & Armstrong, 2015). The Cole-Moore shift is also seen in the gating currents underlying the conductance (F. Bezanilla et al., 1994; Francisco Bezanilla et al., 1982). The Cole-Moore shift can be elicited in several ways. It is typically measured following a prepulse of increasing hyperpolarized voltage before a depolarization (Figure 7A, top left) or increasingly long hyperpolarizing prepulses (Figure 7A bottom left). The classical explanation for the Cole-Moore shift is that a hyperpolarization populates closed states that are further removed from the open state, thus delaying the opening as the channel has to traverse more states before opening (Cole & Moore, 1960). However, the molecular basis of the Cole-Moore shift remains unknown (Hoshi & Armstrong, 2015).

**Figure 7.**
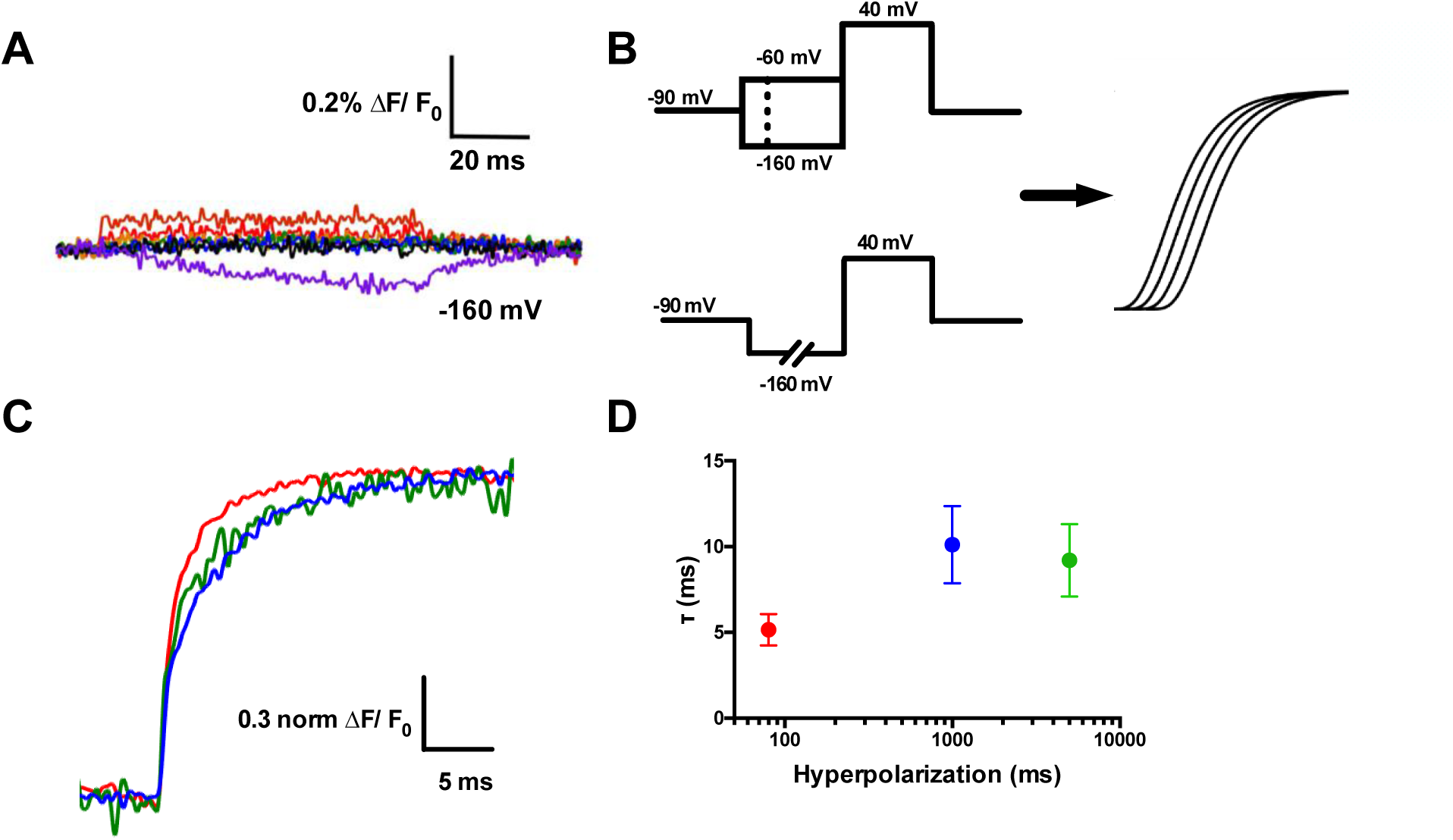
R1C-qBBr interaction with F290 provides the basis of the Cole-Moore shift. (A) A family of fluorescence traces for R1C-qBBr:W454A;F290W from a pulse protocol as in (6A). Note the slow fluorescence response when hyperpolarized to −160 mV (purple) and the residual signals during depolarizations. (B) Two pulse protocols that induce a Cole-Moore shift: a variable voltage pulse before a depolarization (top) or a prolonged hyperpolarization pulse before depolarization (bottom) and a representation of the resulting Cole-Moore shifts of the ionic currents. (C) Representative normalized fluorescence traces for R1C-qBBr:W454A;F290W using the pulse protocol from (B, bottom),with a −160 mV prepulse with a variable duration for 80 ms (red), 1 s (blue), and 5 s (green). (D) Comparison of the **τ**_weighted_ of the change in qBBr fluorescence in (C). (80 ms pulses, n=4; 1 s pulses, n=5; 5 s pulses, n=4). Data are shown as mean ± SEM.

While measuring R1C-qBBr:W454A;F290W fluorescence, we observed no appreciable fluorescence signal above the residual fluorescence in response to depolarizing pulses from −120 mV (Figure 7A). However, in response to a hyperpolarizing pulse to −160 mV from −120 mV, we observed a small, markedly slow reduction in fluorescence (Figure 7A). This fluorescence change did not have the same kinetics of a small measurable gating current for the same pulse protocol (Figure 7— figure supplement 1). When a protocol used to induce and measure a Cole-Moore shift (Figure 7B, left bottom) was applied, the R1C-qBBr:W454A;F290W fluorescence response was larger than the residual response seen in Fig. 7A, and it became slower as the duration of the hyperpolarizing pulse increased (Figure 7C), as expected if this fluorescence is associated with the Cole-Moore shift of the channel.

Previous studies have shown that at negative potentials R1C in the closed state can spontaneously link to I287C, which is a full turn above F290 (Campos et al., 2007). Here, we observe that at extreme hyperpolarizing potentials (−160 mV) R1C-qBBr moves near F290W, producing a slow quenching of fluorescence. We explain this as a result of moving the VSD with a strong hyperpolarization to such an extreme intracellular position that we populate other closed states of the voltage sensor that are responsible for the Cole-Moore shift. These closed states are tracked by the fluorescence change produced by an interaction between F290W and R1C-qBBr. A similar rarely observed closed state predicted by metal-ion bridges, in which R1 transitions from a position extracellular to F290 to a position intracellular to F290, was also suggested to provide a potential explanation of the Cole-Moore shift (Henrion et al., 2012).

### Voltage-sensitive membrane protein CiVSP is interrogable by qBBr

Our technique presented here allows for the interrogation of other VSD movements. Another excellent candidate for qBBr mapping is CiVSP. With the reported crystal structure (Li et al., 2014) we were able to begin qBBr mapping of this VSD (Figure 8A). Using the CiVSP R217Q background (Dimitrov et al., 2007) we mutated the first gating charge, R223, to a cysteine and labeled it with qBBr. The CiVSP R1C-qBBr showed voltage-dependent fluorescence changes (Figure 8B, top). As with Shaker R1C-qBBr we were able to identify the source of the fluorescent change. By mutating tyrosine Y206 to an alanine, we abolished the voltage dependent fluorescence (Figure 8B, bottom), while the voltage dependence of CiVSP with the Y206A mutation remained intact (Figure 8C). This result is a starting point for investigating the movement of the gating charges of this VSD.

**Figure 8.**
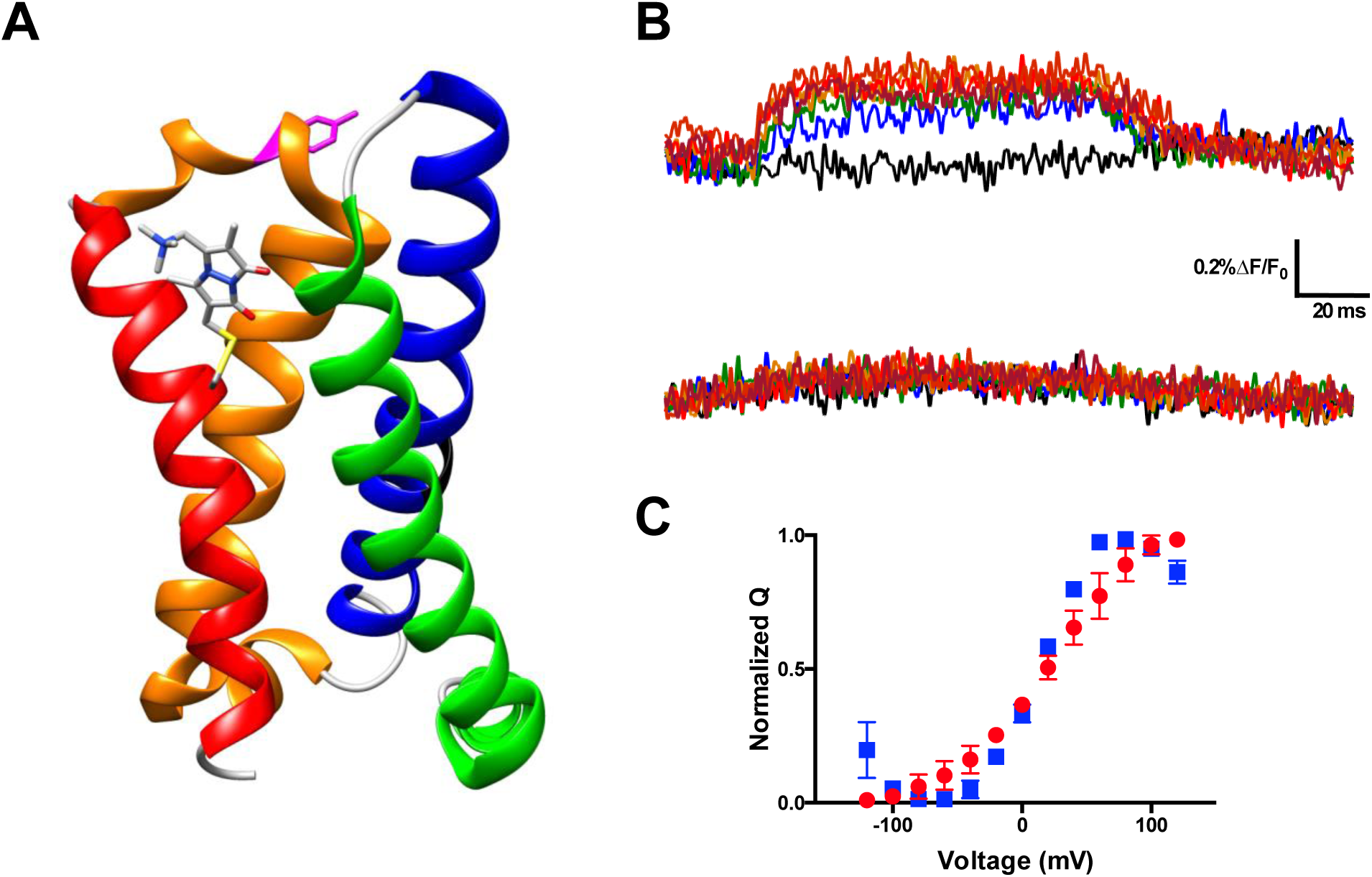
Voltage-sensitive membrane protein CiVSP is interrogable by qBBr. (A) CiVSP structure (PDB: 4G7V) with qBBr attached at R1C and highlighting residue Y206. Transmembrane domains (S1-S4) are colored with S1, green; S2, blue; S3, orange; S4, red. (B) Representative fluorescence traces of CiVSP R217Q R1C-qBBr (top) and CiVSP R217Q R1C-qBBr:Y206A (bottom) (C) Normalized QV curves comparing CiVSP R217Q R1C-qBBr (red circles, n=4) and CiVSP R217Q R1C-qBBr:Y206A (blue squares, n=4). Data are shown as mean ± SEM.

## Discussion

In this paper we have demonstrated a technique that allows tracking of a gating charge surrogate in the pathway of a voltage sensor using qBBr. We have obtained new information on the trajectories of the first two gating charges in Shaker (Figures 4 and 5). We propose that this is a flexible tool that should prove readily applicable to other voltage-sensitive proteins. As we discuss the details of the trajectories, we will point out some of the strengths and limitations of the technique.

### Spatial resolution

In principle, one would expect qBBr mapping to be limited in its spatial resolution. Tryptophan-induced quenching (TrIQ) and tyrosine induced quenching (TyrIQ) of bimane dyes have been proposed as methods for measuring molecular distances (Brunette & Farrens, 2014; Mansoor et al., 2002, 2010). However, the maximum alpha carbon to alpha carbon distance over which TrIQ occurs with qBBr has been found to be around 10 to 11 Å (Mansoor et al., 2010), or possibly as high as 15 Å (Brunette & Farrens, 2014). It is clear that measurements of static distances with this method would have poor spatial resolution, and it would seem that quenching over such long distances would make it very difficult to trace any detailed pathway of qBBr. However, because we are measuring a change in fluorescence as a function of time as the qBBr evolves in its pathway, we obtain much higher spatial resolution than measuring static fluorescence quenching in two positions. This is because we can detect very small changes in relative fluorescence over time as the qBBr approaches or recedes from the quencher.

Take as an example qBBr moving from a position adjacent to W to a distance of 7 Å away from the quencher during a depolarization. The qBBr-W distance is always shorter than the 10 Å known to be the qBBr-W quenching distance. However, the degree of quenching will vary from extremely high when qBBr is close to W to less as it moves to its final position. The change in fluorescence may be only 1 to 2%, but this change still reveals that the qBBr-quencher distance is increasing. In other words, we infer that it moved away even though we do not know the exact distance.

We cannot absolutely calibrate the degree of quenching, mainly because of spurious background fluorescence. Therefore, the technique presented here takes full advantage of the insertion of a quencher in the putative path: when a fluorescence change occurs during a voltage pulse, we can infer the charge is moving with respect to the quencher while if the signal does not change, it means that either the quencher is far from the charge’s path or there is no movement with respect to the quencher.

As a result, even though the interpretation is qualitative rather than quantitative, our method not only constrains the resting and active positions of a particular gating charge but also, by collecting data from many locations, generates a trajectory of the transition pathway of the gating charge.

### Kinetics of trajectories are different for each charge and depend on initial conditions

As laid out in equation 2, in dynamic qBBr-W quenching, a two-state model predicts that the kinetics of the fluorescence is independent of the position of qBBr with respect to the quencher. If qBBr moves between several discrete positions, then kinetics of the fluorescence is affected by the relative distance with respect to the quencher (Equation 3). In a 3-state model the general prediction is a time course of fluorescence with increases and decreases during the voltage pulse that moves the charge. However, when the dwell time of the intermediate state is brief, then the fluorescence change is monotonic and has two time constants. In all our studies, we do not see increases and decreases of fluorescence during a single pulse. Instead, fluorescence changes are always in the same direction (either an increase or a decrease), albeit with different kinetics and more than one time constant. The different kinetics observed between different constructs, therefore, may be explained by the brief dwell time in the intermediate state; however, we cannot exclude that the introduction of a W mutation alters the underlying gating current kinetics. Consequently, although it would be ideal to be able to compare kinetics of one construct to another, the possible differences produced by the W mutations preclude us from doing so.

However, using the same construct, voltage pulses of different durations or potential can be used to interrogate different transitions of the voltage sensor, and the kinetics of different processes from the same construct can then be compared. For example, the deactivation kinetics of the fluorescence change produced by R1C-qBBr:W454A;T326W are more rapid than its activation kinetics, suggesting that the interaction of R1C-qBBr with T326W is different during activation than during deactivation. Additionally, we observe that the QV and FVs of deactivation have a leftward shift to that of activation, thus demonstrating that the path of deactivation is different than that of activation. This in turn informs us that the deactivation pathway of R1C-qBBr must be different from its activation pathway, a conclusion that would be difficult to obtain with any other method. Future studies could use qBBr optical mapping to investigate the movement of discrete gating charges during conductive events to correlate features of gating with ion conductance.

Another example is the slow R1C-qBBr:W454A;F290W fluorescence change upon hyperpolarization. The kinetics of this interaction are markedly different from the kinetics of the main motion of the voltage sensor. This result gives direct evidence that R1 can populate a region of the sensor closer to the intracellular side when the membrane is strongly hyperpolarized and provides a molecular basis for the Cole-Moore shift.

### Trajectories of discrete gating charges

Our data indicate that in the normal resting state, it appears that R1 does not come into close contact with F290 or I294 in the S2 segment, while R2 does come into proximity with I294. In the normal active state, R1 comes into close contact with W454 and Y415, while R2 comes into close contact with Y415, but does not interact with W454. One observation potentially related to the position of the resting state is that two of our tested constructs were lethal: R1C:W454A;I287W and R2C:F290W. Whether the lethality of these constructs stems from a disruption of the normal resting state interactions of discrete gating charges with the gating pore remains to be investigated. However, R1 closely interacting with I287 in the resting state is in good agreement with earlier findings using disulfide bonding (Campos et al., 2007), fluorimetry (Pathak et al., 2007), and modeling (Henrion et al., 2012; Vargas et al., 2011).

The ability to optically track a gating charge in real time is a powerful technique, despite its limitations. It allows for the combination of crystallographic studies and functional data to understand the movement of the VSD. In general, our data with R1 and R2 supports a sliding helix model of movement of the gating charges but adds a more detailed view of the movement. Each discrete gating charge undergoes both a rotation with translation with an angle of about 60° with respect to the plane of the membrane. Interestingly, the rotation of the R1 appears greater than the rotation of the R2, suggesting that some of the movement may be coming from the movement of the side chain, or a transient change of the helical conformation from an α helix to a 3_10_ helix (Bassetto et al., 2020; Chakrapani et al., 2010; Henrion et al., 2012; Schwaiger et al., 2011) rather than exclusively from the movement of the voltage sensor backbone. We expect that this work can be expanded with molecular modeling. We may be able to test whether the charges move independently of each other, with the first and second gating charge moving first and then tugging the rest of the VSD along via the backbone of the S4 segment (Horng et al., 2019).

The qBBr fluorescence data constrains the normal resting state of R1 to a position extracellular to F290 and indicate that interactions between this residue and R1 may be responsible for the Cole-Moore shift by moving the first charge deeper into the intracellular side. This result is consistent with the shallow Q-V curve recorded at very negative potentials and also with the fact that the ‘closed state’ is in fact a collection of closed states in agreement with the increase in entropy when evolving from the open to the closed state (Claydon et al., 2007; Rodríguez et al., 1998).

### Extension to other voltage sensors and gating charges

Numerous voltage sensors have been examined with site-directed fluorimetry: Shaker (Cha & Bezanilla, 1997; Mannuzzu et al., 1996), other Kv1-type channels (Peters et al., 2009; Vaid et al., 2008), Kv7 (Kim et al., 2017; Osteen et al., 2010; Ruscic et al., 2013), Kv10 and Kv11 (Schönherr et al., 2002; Smith & Yellen, 2002), Nav (Cha et al., 1999; Chanda & Bezanilla, 2002; Wang et al., 2016), voltage-gated calcium channels (Pantazis et al., 2014), BK (Savalli et al., 2006), HCN (Bruening-Wright & Larsson, 2007), Catsper (Arima et al., 2018), and CiVSP (Kohout et al., 2008; Villalba-Galea et al., 2008). Extension of the qBBr WY-induced quenching technique to other voltage sensors is expected to provide new findings because this technique traces the actual charge trajectories during voltage sensing. As a proof-of-principle, we demonstrated qBBr can be applied to CiVSP, an evolutionarily distant voltage sensor. Interestingly, we uncovered an interaction between R1 of CiVSP and a tyrosine residue in the S3-S4 linker that is in good agreement with a reported crystal structure of the voltage-sensing domain in the active state (Li et al., 2014). Further mapping of additional voltage sensor pathways with the qBBr method described here should improve our understanding of how voltage sensors and their discrete gating charges move and provide new insight into where the electric field is focused, which side charges interact with the VSD, and the differences in gating charge trajectories. Finally, a present limitation of the technique is that qBBr mapping of gating charges is limited by labeling accessibility to gating charge residues mutated to cysteine. For example, we were unable to link qBBr to the third most extracellular Shaker gating charge (R368). Future experiments using unnatural amino acid versions of qBBr or other fluorescent amino acids with characteristics similar to qBBr (Leisle et al., 2015) may allow for the expansion of this technique to additional charges in the voltage sensor.

## Materials and Methods

### Generation of constructs

Mutations of *Shaker* or *CiVSP* DNA were made on the *Shaker* Δ6-46 W434F background (Perozo et al., 1993) or the *CiVSP* C363S background (Murata et al., 2005), respectively, using the QuikChange II site directed mutagenesis kit (Agilent, Santa Clara, CA) with primers purchased from Integrated DNA Technologies. *Shaker* DNA was linearized using NotI (New England Biolabs, Ipswich, MA) and *CiVSP* DNA was linearized using XbaI (New England Biolabs); both were cleaned up with a NucleoSpin Gel and PCR Clean-up kit (Macherey-Nagel, Bethlehem, PA). *Shaker* and *CiVSP* cRNAs were then synthesized from linearized DNA using the mMESSAGE mMACHINE T7 or SP6 transcription kit, respectively (Life Technologies, Carlsbad, CA).

### Oocyte preparation

Oocytes were harvested from *Xenopus laevis* in accordance with experimental protocols approved by the University of Chicago Animal Care and Use Committee. Unless specified, all chemicals were obtained from Sigma-Aldrich (St. Louis, MO). Following collagenase digestion of the follicular membrane, oocytes were maintained in standard oocyte solution containing 96 mM NaCl, 2 mM KCl, 1 mM MgCl_2_, 1.8 mM CaCl_2_, 10 mM HEPES, and 50 μg/ml of gentamicin, set with NaOH to pH 7.4, for up to 36 hours prior to injection with 50 ng of cRNA. Following cRNA injection, oocytes were incubated at 16°C in standard oocyte solution for three to six days prior to recording.

### Labeling with qBBr

The labeling solution consisted of depolarizing solution comprised of 120 mM KCl, 2 mM CaCl_2_ and 10 mM HEPES at pH 7.4 with 1-2 mM qBBr (Sigma-Aldrich or Toronto Research Chemicals, North York, ON, Canada) added fresh to the solution upon each preparation. Oocytes were maintained in the solution for at least fifteen minutes and removed and washed in standard oocyte solution 5-20 minutes before recordings on them were performed. At the longest, oocytes were maintained in labeling solution for 120 minutes; however, due to the positive charge and impermeability of qBBr, no differences were observed between oocytes labeled for long durations versus short durations.

### Simultaneous fluorescence and electrophysiological recordings

Simultaneous electrophysiological and fluorescence recordings were performed at room temperature using the cut-open oocyte voltage-clamp technique, similar to prior descriptions (Cha & Bezanilla, 1997; Lacroix et al., 2012; Villalba-Galea et al., 2008). External solution contained 115 mM N-methyl-D-glucamine, 2 mM Ca(OH)_2_, and 20 mM HEPES taken to a pH between 7.4 and 7.5 with methanesulfonic acid. Internal solution was like external solution, but with 2 mM EGTA in place of Ca(OH)_2_. Pipettes were pulled at a resistance of 0.2 to 1.0 MΩ and filled with 3M KCl. Excitation was performed with a mounted 420 nm LED (ThorLabs, Newton, NJ) reflected by a 455 nm long-pass dichroic (Chroma, Bellows Falls, VT) through a 40X water-immersion objective (LUMPlan FL N, Olympus, Center Valley, PA); emission was collected through the dichroic and a 475 nm long-pass filter (Chroma). Emission was integrated over each sampling period through a home-built integrator, collected by a PIN-020A photodiode (UDT Technologies, Torrance, CA), and amplified by a patch-clamp amplifier (L/M-EPC-7, LIST Medical Electronics, Darmstadt, West Germany). Voltage-clamp and electrical measurements were performed with a CA-1B amplifier (Dagan, Minneapolis, MN). The LED and voltage-clamp were controlled through Gpatch, an in-house acquisition program, and an SB6711-A4D4 board (Innovative Integration, Simi Valley, CA). Recordings at each voltage step were an average of four traces, taken consecutively, sampled at 50-100 kHz and filtered at 5-10 kHz.

### Data analysis

Recordings were filtered and analyzed offline using custom MATLAB (Mathworks, Natick, MA) scripts (Treger et al., 2015). Filtering of fluorescence for display and analysis was performed using a digitized Bessel filter with a cutoff frequency between 500 Hz and 1 kHz. ΔF/F_0_ was calculated following a linear baseline subtraction of the period 6 to 1.5 s prior to the electrical potential change of interest. Charge was calculated from gating currents following linear baseline subtraction of user-defined periods based on when gating current amplitudes returned to zero. Time constant measurements of kinetics of both fluorescence and gating were calculated using either a single exponential fit or a weighted double exponential fit to the rising or falling phase of the fluorescence as appropriate, or to the decay component of the gating current.

Images of the Shaker and CiVSP proteins with qBBr and tryptophan substitutions were created using UCSF Chimera (Pettersen et al., 2004).

### Quantification and statistical analysis

Statistical details of experiments can be found in figure legends. N represents the number of oocytes, or biological replicates, and data are shown as mean ± SEM. Calculations were performed in Prism (GraphPad, La Jolla, CA).

## Supplemental Figures

**Figure 2—figure supplement 1.**
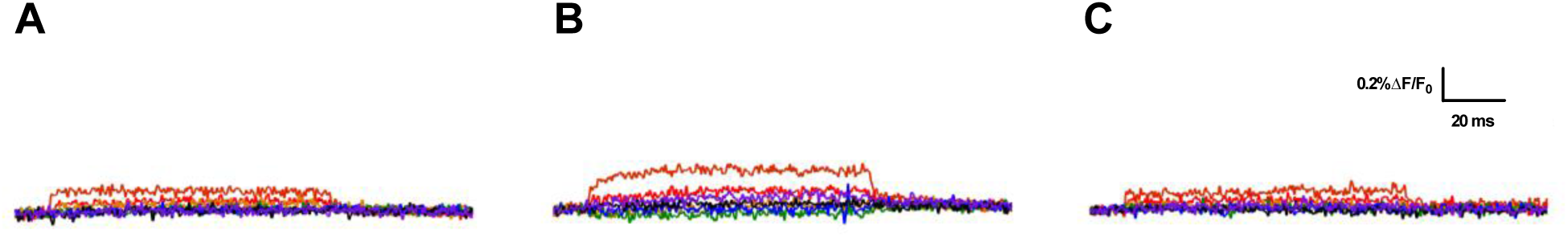
R1C-qBBr:W454A and R2C-qBBr fluorescence signals are not produced by endogenous tryptophans or tyrosines. Representative fluorescence traces of qBBr labeled (A) R1C-qBBr:W454A;W289F (B) R1C-qBBr:W454A;Y323F (C) R2C-qBBr W289F;Y323F. In all panels, membrane potentials during the pulse are: brown, +80 mV; red, +40 mV; orange, 0 mV; green, −40 mV; blue, −80 mV; black, −120 mV; purple, −160 mV.

**Figure 6—figure supplement 1.**
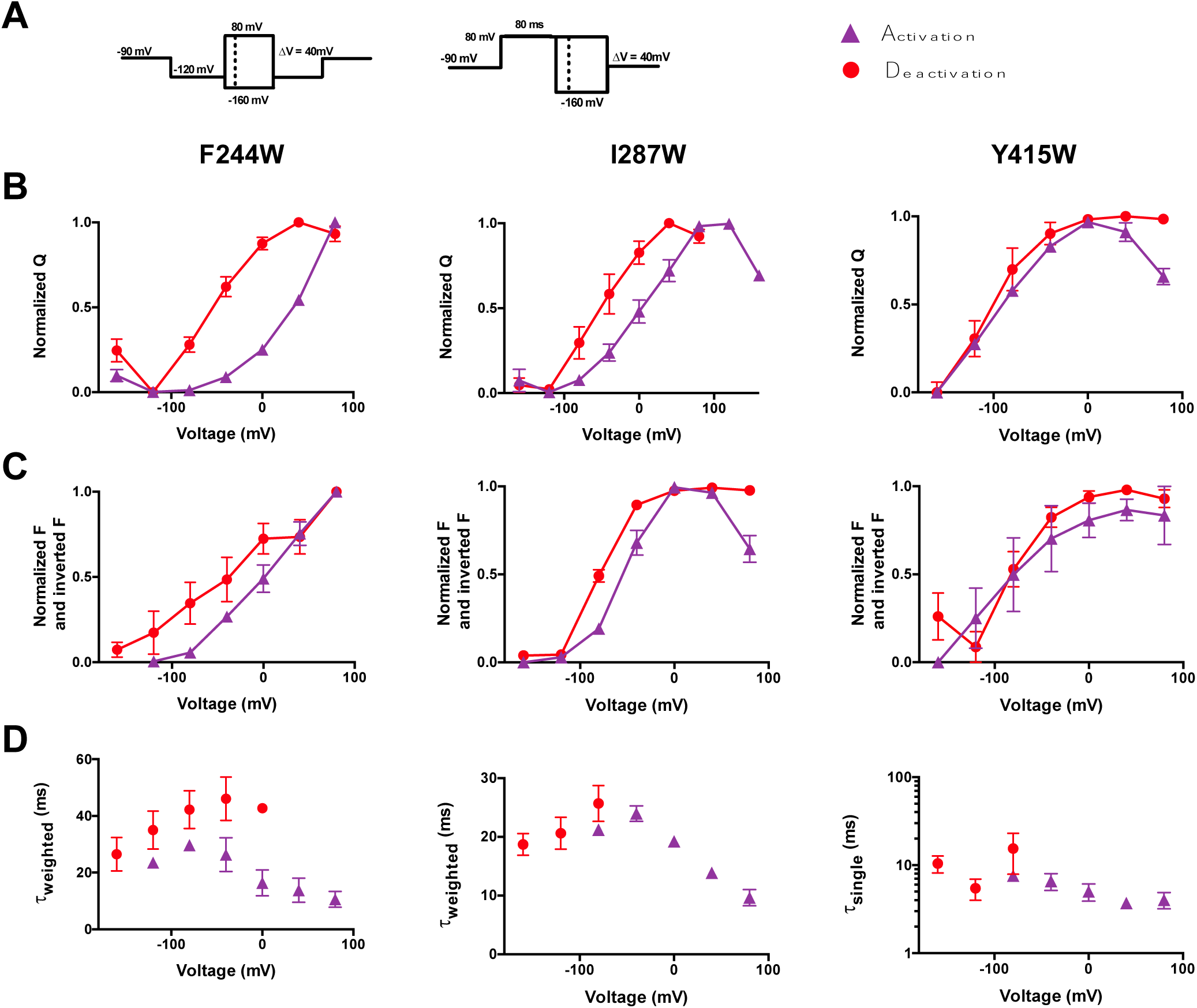
The VSD deactivation path of R2C-qBBr differs from that of activation. (A) Activation (purple) and deactivation (red) protocols. Shading indicates where gating and fluorescence were measured for (B-D). (B) Comparison of activation (purple triangles) and deactivation (red circles) QVs for R2C-qBBr:F244W (left, n=4), R2C-qBBr:I287W (center, n=5), and R2C-qBBr:Y415W (right, n=4). (C) As in (B), but a comparison of activation and deactivation FVs, rather than QVs. (D) Comparison of single exponential **τ**_act_ (red circles, n=4) to **τ**_dea_ (purple triangles, n=7) of R2C-qBBr:F244W (left), weighted **τ**_act_ (red circles, n=4) to **τ**_dea_ (purple triangles, n=5) of R2C-qBBr:I287W (center), and weighted **τ**_act_ (red circles, n=5) to **τ**_dea_ (purple triangles, n=4) of R2C-qBBr:Y415W (right). Data are shown as mean ± SEM.

**Figure 6—figure supplement 2.**
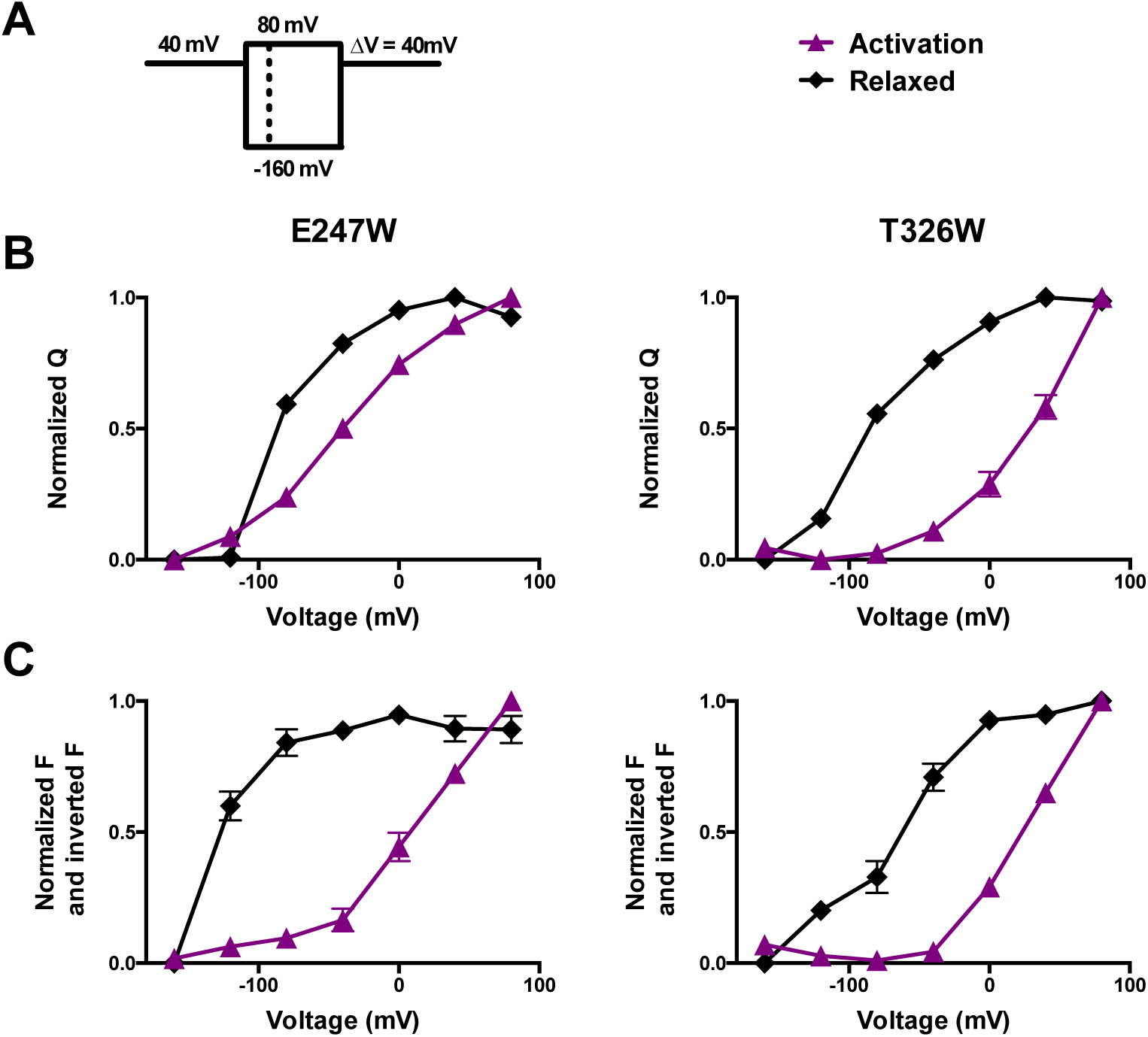
R1C-qBBr fluorescence visualizes the relaxed state. (A) A relaxation pulse protocol. (B) Comparison of activation (purple, triangles) and derelaxation (black, diamonds) QVs for R1C-qBBr:W454A;E247W (left, n=4) and R1C-qBBr:W454A;T326W (right, n=4). (C) Inverted FV_normalized_ and FV_normalized_ for activation and derelaxation for R1C-qBBr:W454A;E247W (left, activation n=8, derelaxation n=6) and R1C-qBBr:W454A;T326W (right, activation n=5, derelaxation n=6). Data are shown as mean ± SEM.

**Figure 7—figure supplement 1.**
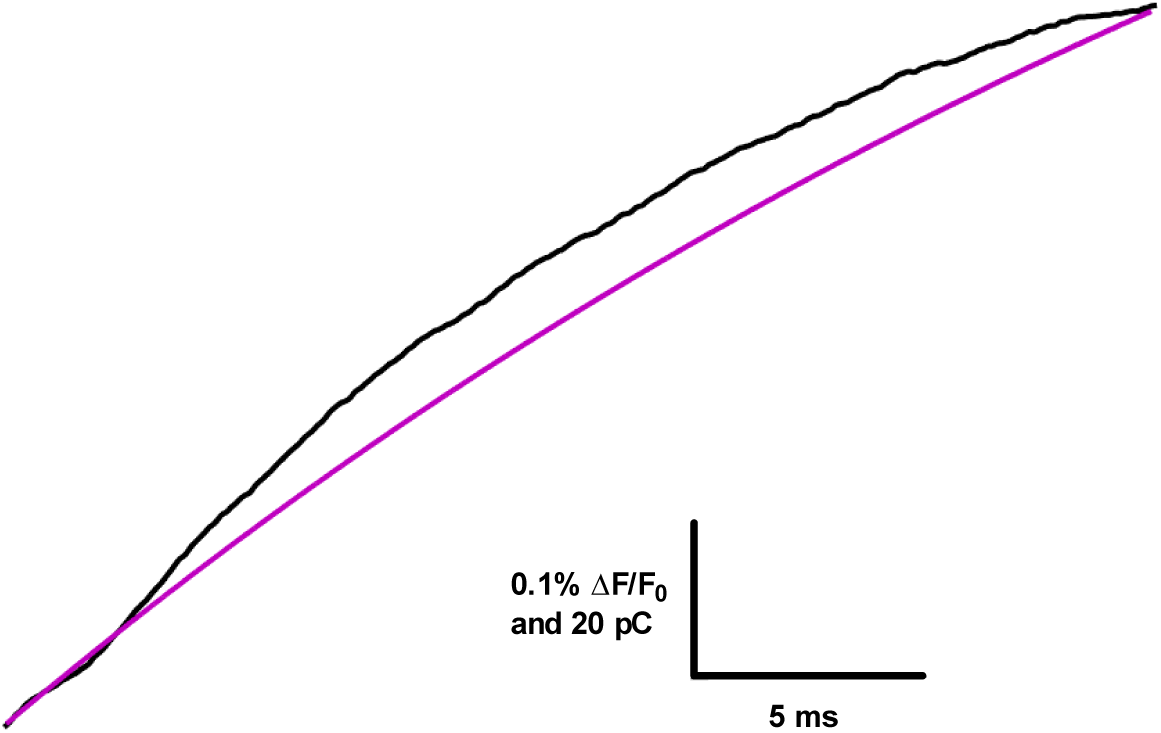
R1C-qBBr:W454A;F290W hyperpolarization-induced gating charge and fluorescence signal time course. Data is taken from a pulse from −120 mV to −160 mV. The integrated gating current of all four charges moving through the electric field (gating charge, Q, black) is more rapid than the hyperpolarization-induced fluorescence signal of R1C-qBBr:W454A;F290W, shown as the single exponential fit to the inverted fluorescence signal (purple). The fluorescence signal only follows the movement of the first charge. Traces are aligned to each other, but the fluorescence signal is still slower than the gating charge.

